# Polyhydroxyalkanoate granule accumulation makes optical density measurement an unreliable method for bacterial growth assessment in *Burkholderia thailandensis*

**DOI:** 10.1101/682161

**Authors:** Sarah Martinez, Eric Déziel

## Abstract

Optical density (OD) measurement is the standard method used in microbiology for estimating bacterial concentrations in cultures. However, most studies do not compare these measurements with viable cell counts and assume that they reflect the real cell concentration. *Burkholderia thailandensis* was recently identified as a polyhydroxyalkanoate (PHA) producer. PHA biosynthesis seems to be coded by an ortholog of the *Cupriavidus necator phaC* gene. When growing cultures of wildtype strain E264 and an isogenic *phaC*- mutant, we noted a difference in their OD_600_ values, although viable cell counts indicated similar growth. Investigating the cellular morphologies of both strains, we found that under our conditions the wildtype strain was full of PHA granules, deforming the cells, while the mutant contained no granules. These factors apparently affected the light scattering, making the OD_600_ values no longer representative of cell density. We show a direct correlation between OD_600_ values and the accumulation of PHA. We conclude that OD measurement is unreliable for growth evaluation of *B. thailandensis* because of PHA production. This study also suggests that *B. thailandensis* could represent an excellent candidate for PHA bioproduction. Correlation between OD measurements and viable cell counts should be verified on any study realized in *B. thailandensis*.

---

The soil saprophyte *Burkholderia thailandensis* was recently identified as a producer of polyhydroxyalkanoates (PHAs) (Funston, Tsaousi et al. 2017) and then studied for the co-production of PHAs and rhamnolipid in used cook oil (Kourmentza, Costa et al. 2018). PHAs are polymers of hydroxylated fatty acids that are synthesized by different prokaryotic microorganisms as intracellular carbon storage material (Anderson and Dawes 1990). PHA granules are produced when carbon is present in excess combined with a nitrogen or phosphate limitation (Poblete-Castro, Escapa et al. 2012).

PHA synthases and their genetic organisation are well characterized (Rehm and Steinbüchel 1999), with two classes of PHAs principally documented: class 1 for short chain length-PHAs (scl-PHAs), products of *phaC-phaA-phaB* gene clusters featured in the *Ralstonia*/*Cupriavidus* genus and class 2 for medium chain length PHAs (mcl-PHAs), products of *phaC1-phaZ-phaC2* gene clusters typical of the *Pseudomonas* genus.

PHA synthases are encoded by the *phaC* homologues (Solaiman and Ashby 2005). Investigating the genome of the prototypical *B. thailandensis* strain E264, (Funston, Tsaousi et al. 2017) reported that the BTH_I2255 gene shows similarity with *Pseudomonas*-type poly-(3-hydroxyalkanoate) polymerases coded by *phaC1* and *phaC2* with a percentage of identity of 40 and 39% respectively. Actually, we rather found a 75% identity between BTH_I2255 and the ortholog of the *C. necator* PHA biosynthesis gene *phaC* (NC_015726.1). Furthermore, the genomic context of this gene reveals the immediate proximity of *phaA* and *phaB* orthologs, coding an acetyl-CoA acetyltransferase and an acetoacetyl-CoA reductase, responsible for the precursors biosynthesis, with 82% and 83% identity respectively with the corresponding *C. necator* genes (CAJ92573.1 and CAJ92574.1), suggesting that a complete class 1 PHA biosynthesis machinery is present in *B. thailandensis* for the production of scl-PHAs. This is compatible with the affiliation of both *Burkholderia* and *Cupriavidus* in the class β-Proteobacteria. We indeed also found an ortholog of the *Pseudomonas aeruginosa* mcl-PHA biosynthesis gene, *phaC1* (NC_002516.2). However, it corresponds to locus tag BTH_II0418, with 74% identity. No *P. aeruginosa phaZ* gene ortholog was found, indicating that a mcl-PHAs biosynthesis is not complete in *B. thailandensis*.

Optical density (OD) measurements at 600 nm are the standard approach used in microbiology labs for estimating the bacterial concentration in a liquid culture (Stevenson, McVey et al. 2016). With homogenous cultures, it is generally found that OD remains proportional to bacterial density throughout the positive phases of growth of the cultures (Monod 1942). When this requirement is fulfilled, OD determinations can provide an adequate and extremely convenient method of estimating bacterial density. However, these growth estimations can be misleading. Indeed, there are relatively few studies comparing these estimations with viable cell counts. Optical density is a measure of the light scattered by the bacteria present in the suspension. Indeed, in a non-scattering sample, the attenuation in light transmission between the light source and the detector is caused by the absorbance of light by the sample. But, in a scattering sample, such as a bacterial suspension, the light reaching the detector is further reduced by the scattering of light. This decrease creates the illusion of an increase in sample absorbance, while it is actually a measure of turbidity (Matlock, Beringer et al. 2011).

We demonstrate here that PHAs accumulated as intracellular granules in *B. thailandensis* cells affect the size and shape of the bacteria sufficiently to increase the possibility of multiple light scattering events, making OD_600_ values unreliable for growth evaluation.

## Optical density at 600 nm is not representative for growth measurement in B. thailandensis

Wildtype *B. thailandensis* E264 (WT), and isogenic *phaC*- and *phaC1*- mutants obtained from the transposon mutant library (Gallagher, Ramage et al. 2013), were grown in NB medium complemented with 2% glycerol. OD_600_ and colony forming unit (CFU) counts were monitored during 4 days. Interestingly, OD_600_ measurements indicated that the *phaC*^−^ mutant reached lower values than the WT strain (Fig. 1a). On the other hand, loss of *phaC1* did not affect OD_600_ values. This was suggesting that the *phaC*^−^ mutant has a growth defect. Unexpectedly, CFU counts revealed the concentration of bacteria present was similar for all three strains (Fig. 1b). Thus, there is a lack of correlation between OD measures and cell density.

**Figure 1:**
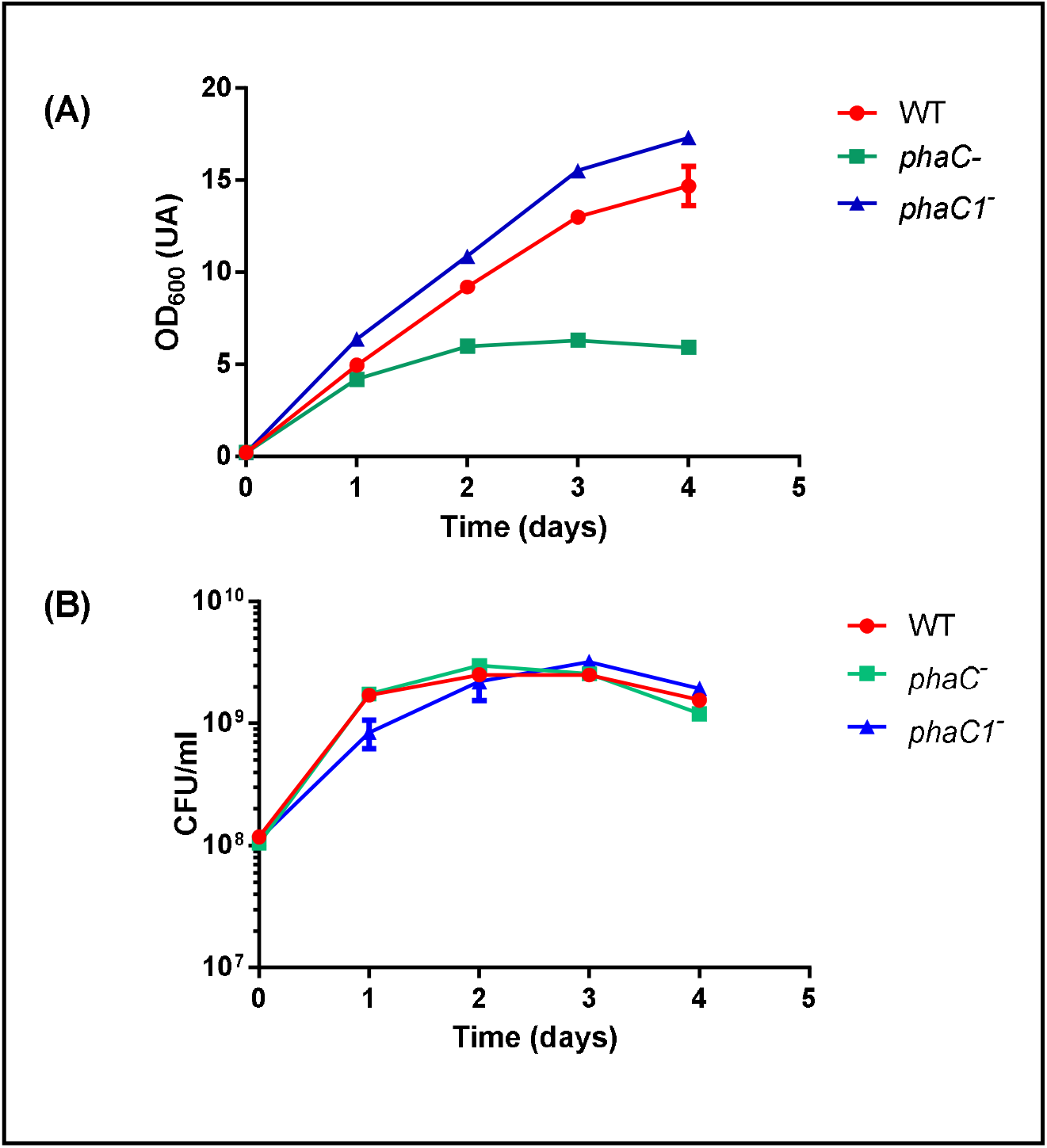
Optical densities at 600 nm and CFUs may not be correlated in *B. thailandensis* cultures. The error bars represent standard deviation from the mean (*n* = 3 independent cultures). E264 and *phaC*- and *phaC1*- mutants were grown in 3 mL Nutrient broth (NB; Difco) medium supplemented with 2% (w/v) glycerol. The cultures were incubated at 30°C with rotation at 240 rpm in a TC-7 roller drum (New Brunswick, Canada) for 16h. These seed cultures were then diluted in flasks containing 25 mL of NB medium supplemented with 2% (w/v) glycerol at OD_600_ = 0.1 and then incubated at 30°C with agitation (200 rpm) in a gyratory shaker (Infors) for 5 days. Samples were collected daily and bacterial growth was estimated by OD_600_ measurements (A) and by viable counts (B), determined by plating serial dilutions on LB agar plates, incubated for 24 hours at 37°C. Colonies were counted to determine bacterial cell count (CFU/mL) for each time point.

We identified the *phaC* gene as responsible for the PHA production in *B. thailandensis*. This is consistent with the PHA composition described in a recent study, where scl-PHA, also named polyhydoxybutyrate (PHB), were characterized (Kourmentza, Costa et al. 2018).

## PHA production is responsible for the OD difference between the wildtype strain and phaC- mutant

During cultivation, PHA production was verified for the three strains. Two strategies were employed. First, PHA biosynthesis was evaluated by Nile blue staining and fluorescence quantification. Indeed, Nile blue is a fluorescent dye specific for PHA granules (Page and Tenove 1996). Nile Blue staining relies on a linear correlation obtained between intracellular PHA concentrations and the corresponding fluorescence intensities (Zuriani, Vigneswari et al. 2013). PHA accumulation was measured as described by (Oshiki, Satoh et al. 2011), with some adjustments. Briefly, samples were collected and submitted to a heat choc. Addition of Nile blue in the lysate allowed to evaluate PHA production by measuring emitted fluorescence. Data show that the *phaC* mutant does not produce PHAs as no fluorescence was detected (Fig. 2A). In contrast the WT samples showed increasing fluorescence values, indicating the presence of PHA granules, as did the *phaC1*- mutant, strongly suggesting that this gene is not involved in PHA production, at least under our conditions.

**Figure 2:**
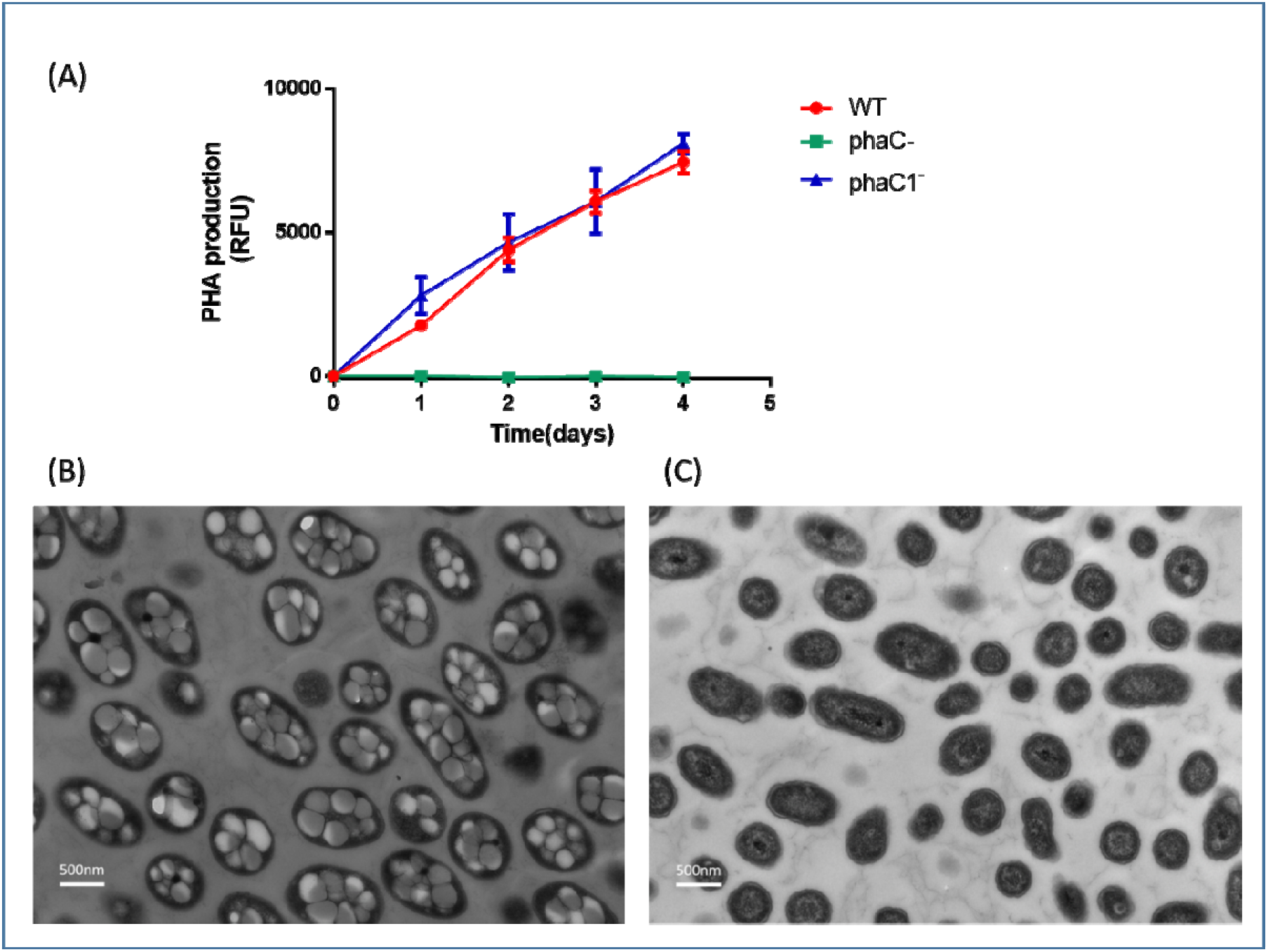
*phaC*^−^ mutant of *B. thailandensis* E264 do not produce PHAs. (A) PHA production for the wild type strain and the *phaC*- and *phaC1*- mutants measured by Nile blue staining. Culture samples of 100 μL were collected at regular time intervals and were centrifuged for 3 min at 10,000 x *g*. Supernatants were discarded and pellets were suspended in 100 μL water. Samples were heated at 100°C for 10 min and transferred on ice for 5 min to lyse the cells. Samples were then centrifuged for 3 min at 10,800 x g, supernatants discarded and pellets suspended in 100 μL water. Samples were transferred to a 96-well plate and an equal volume of a 0.02% (w/v) Nile Blue (Sigma-Aldrich) solution was added to each well. After a 4 min incubation, the fluorescence intensity (excitation 490 nm / emission 590 nm) was measured using a Cytation (Biotek) multimode microplate reader. (B) and (C) are TEM images (x10.000) for the wildtype strain and the *phaC*- mutant respectively. After four days of incubation, 1 mL of each culture were collected and transferred into a sterile 1.5 microcentrifuge tube. After centrifugation (3 min, 8,000 x *g*), the supernatant was discarded and cells were fixed with 2.5% gluteraldehyde in phosphate buffer saline and kept at 4°C overnight. Cells were washed three times at room temperature with 200 μL of washing solution (0.05 M cacodylate sodium, 3% sucrose). Staining was performed with 1.3% osmium tetroxide in collidine buffer. Fixed stained material was progressively dehydrated with increasing concentrations of acetone (25-95%). The material was kept overnight in a 1:1 volume mixture of Spurr resin and acetone, then immersed twice each 2 hours, in a bath of Spurr mixtures. The block containing fixed cells was cut into small pieces, placed in BEEM capsules, filled with Spurr resin and held at room temperature for 16 hours and then polymerized at 60°C for 20-30 hours. Ultrathin sections (70-100 nm thick) were examined by a Hitachi H-7100 electron microscope with an accelerating voltage of 75 kV.

Next, to confirm the data obtained with the Nile blue staining, samples were prepared and observed using transmission electron microscopy (TEM) (Fig. 2B and 2C). Under our culture conditions, the WT strain contained an average of seven big PHA granules/cell, inducing a cell deformation (Fig. 2B). Indeed, WT cells were bulkier than *phaC*- mutant cells, for which TEM confirmed the absence of PHA granules.

Obviously, the shape or size of cells can have an impact on OD measurements (Koch and Ehrenfeld 1968). More recently, the effects of cell size variation was observed with bead suspensions and then confirmed in cultures of *E. coli* and yeast (Stevenson, McVey et al. 2016). A change in cell size throughout the growth curve of an *E. coli* grown under sub-lethal concentrations of ampicillin was demonstrated. This treatment induced a filamentation of the bacteria and thus caused a substantial deviation between OD_600_ and bacterial concentration. During the initial part of the log phase, OD_600_ and bacterial concentration showed the same time dependency but later the cell number remains roughly constant while the OD_600_ increases (Stevenson, McVey et al. 2016). We observed a similar trend here, as when growth seems to have stopped for both the WT and the *phaC*^−^ mutant when viable cell counts are considered, the OD_600_ keeps increasing.

At least under our culture conditions, OD_600_ is actually an indicator of PHA biosynthesis for *B. thailandensis*. Indeed, when PHA production is compared to the OD_600_, the trend line is linear with an R^2^ =0.9845, indicating that PHA biosynthesis is correlated to the OD_600_ values (Fig. 3).

**Figure 3:**
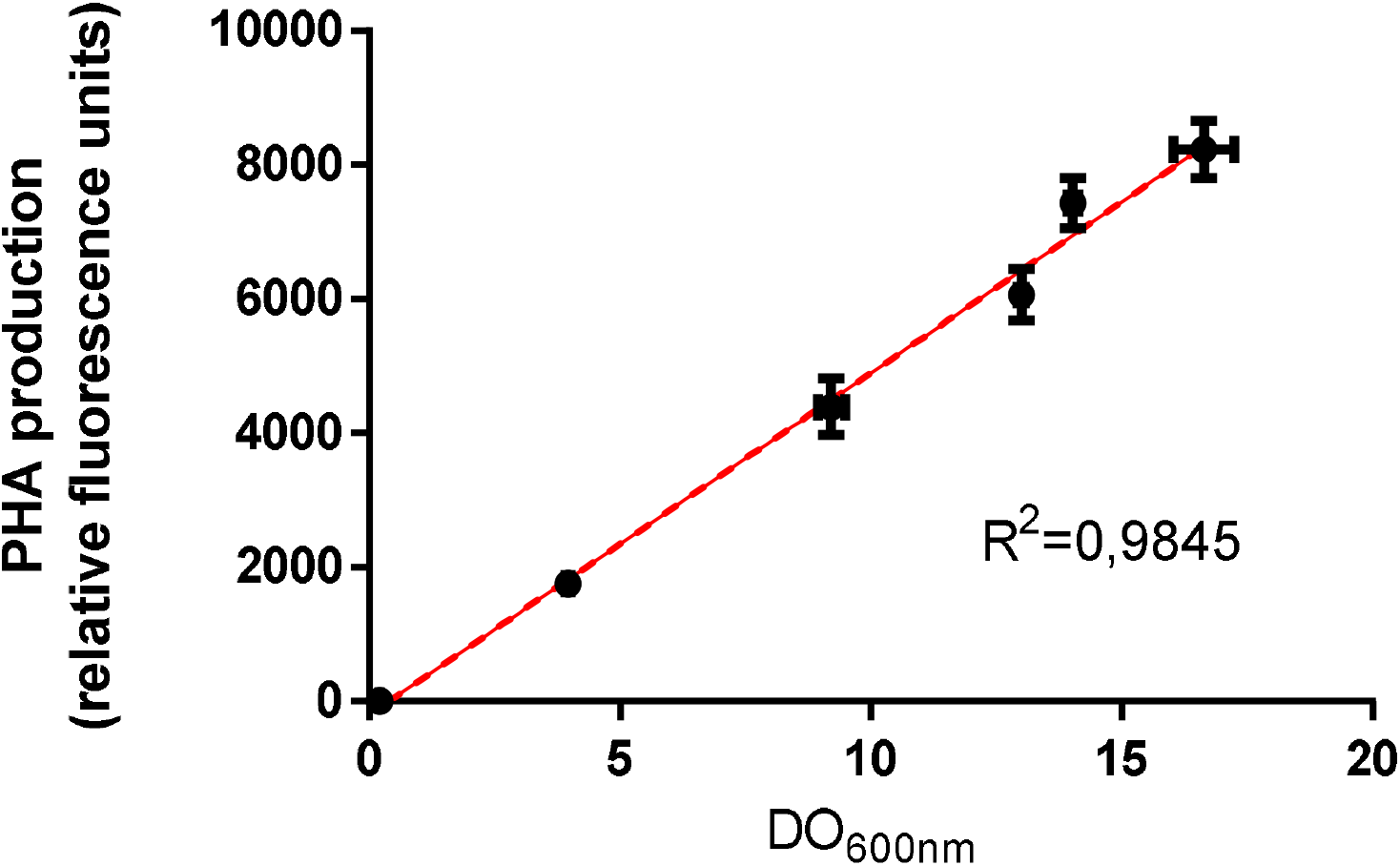
PHA biosynthesis is correlated to OD_600_ measures. PHA biosynthesis is represented depending on the OD_600_ values measured during the growth of the E264 strain in NB medium supplemented with 2% glycerol. The error bars represent standard deviation from the mean (*n* = 3 independent cultures).

In conclusion, we demonstrated that while *B. thailandensis* appears to harbour two distinct PHA biosynthases, only class 1 *phaC* is actually involved in PHA production and not the class 2 *phaC1* homolog. Interestingly, we found here that OD_600_ measurement is not reliable for growth evaluation of *B. thailandensis* when the growth conditions allow PHA production This is consistent with other studies mentioning that turbidimetry is disturbed by the PHA amounts produced. For instance, in *Pseudomonas species*, among which are found well-known PHA producers such as *P. aeruginosa* or *P. putida*, growth measurements in term of viable cells was suggested by using gravimetric methods (Escapa, Del Cerro et al. 2013) since dry cell mass values for growth evaluation have to be adjusted to the PHA content of cells (Escapa, García et al. 2012).

Also, we clearly show that *B. thailandensis* has an excellent potential as a PHA producer, probably comparable to *C. necator* (Mravec, Obruca et al. 2016). PHA production in *B. sacchari* and *B. cepacia* were recently reported, with PHB contents around 88% and 74%, respectively, in a medium containing detoxified wood hydrolyzates (Kucera, Benesova et al. 2017). Furthermore, Kourmentza et *al* (2018) reported that PHB content represents 60% of the dry cell mass when *B. thailandensis* was grown on used cooking oil (Kourmentza, Costa et al. 2018). All together, these observations suggest that studies using optical density measurements for growth assessment might be inconsistent for *Burkholderia* species.

## Acknowledgments

Marie-Christine Groleau for insightful comments and Arnaldo Nakamura for TEM images.

## References

Anderson, A. J. and E. A. Dawes (1990). “Occurrence, metabolism, metabolic role, and industrial uses of bacterial polyhydroxyalkanoates.” Microbiological reviews 54(4): 450–472.

Escapa, I. F., et al. (2013). “The role of GlpR repressor in Pseudomonas putida KT2440 growth and PHA production from glycerol.” Environmental Microbiology 15(1): 93–110.

Escapa, I. F., et al. (2012). “The polyhydroxyalkanoate metabolism controls carbon and energy spillage in Pseudomonas putida.” Environmental Microbiology 14(4): 1049–1063.

Funston, S. J., et al. (2017). “Enhanced rhamnolipid production in Burkholderia thailandensis transposon knockout strains deficient in polyhydroxyalkanoate (PHA) synthesis.” Applied microbiology and biotechnology 101(23-24): 8443–8454.

Gallagher, L. A., et al. (2013). “Sequence-defined transposon mutant library of Burkholderia thailandensis.” MBio 4(6): e00604–00613.

Koch, A. L. and E. Ehrenfeld (1968). “The size and shape of bacteria by light scattering measurements.” Biochimica et Biophysica Acta (BBA)-General Subjects 165(2): 262–273.

Kourmentza, C., et al. (2018). “Burkholderia thailandensis as a microbial cell factory for the bioconversion of used cooking oil to polyhydroxyalkanoates and rhamnolipids.” Bioresource Technology 247: 829–837.

Kucera, D., et al. (2017). “Production of polyhydroxyalkanoates using hydrolyzates of spruce sawdust: Comparison of hydrolyzates detoxification by application of overliming, active carbon, and lignite.” Bioengineering 4(2): 53.

Matlock, B., et al. (2011). “Differences in bacterial optical density measurements between spectrophotometers.” Thermo Scientific Technical Note 52236.

Monod, J. (1942). Recherches sur la croissance des cultures bacteriennes. Sciences naturelles, Université de Paris. PhD: 210.

Mravec, F., et al. (2016). “Accumulation of PHA granules in Cupriavidus necator as seen by confocal fluorescence microscopy.” FEMS Microbiology Letters 363(10): fnw094.

Oshiki, M., et al. (2011). “Rapid quantification of polyhydroxyalkanoates (PHA) concentration in activated sludge with the fluorescent dye Nile blue A.” Water Science and Technology 64(3): 747–753.

Page, W. J. and C. J. Tenove (1996). “Quantitation of poly-β-hydroxybutyrate by fluorescence of bacteria and granules stained with Nile blue A.” Biotechnology Techniques 10(4): 215–220.

Poblete-Castro, I., et al. (2012). “The metabolic response of P. putida KT2442 producing high levels of polyhydroxyalkanoate under single-and multiple-nutrient-limited growth: Highlights from a multi-level omics approach.” Microbial cell factories 11(1): 34.

Rehm, B. H. and A. Steinbüchel (1999). “Biochemical and genetic analysis of PHA synthases and other proteins required for PHA synthesis.” International journal of biological macromolecules 25(1-3): 3–19.

Solaiman, D. K. and R. D. Ashby (2005). “Rapid genetic characterization of poly (hydroxyalkanoate) synthase and its applications.” Biomacromolecules 6(2): 532–537.

Stevenson, K., et al. (2016). “General calibration of microbial growth in microplate readers.” Scientific reports 6: 38828.

Zuriani, R., et al. (2013). “A high throughput Nile red fluorescence method for rapid quantification of intracellular bacterial polyhydroxyalkanoates.” Biotechnology and bioprocess engineering 18(3): 472–478.

